# Connection Failure: Differences in White Matter Microstructure are associated with *5-HTTLPR* but not with Risk-Seeking for Losses

**DOI:** 10.1101/2021.02.03.429606

**Authors:** Philipp T. Neukam, Dirk K. Müller, Yacila I. Deza-Araujo, Shakoor Pooseh, Stephanie H. Witt, Marcella Rietschel, Michael N. Smolka

**Author notes:** Corresponding author, Technische Universität Dresden, Section of Systems Neuroscience, Würzburger Str. 35, 01187 Dresden, Germany.

## Abstract

In a previous study (Neukam, Kroemer et al. 2018), we found for the *5-HTTLPR* genotype higher risk-seeking for losses in S/S vs. L/L carrier, which could not be explained by acutely changed central serotonin levels. This finding alternatively may be the result of reduced top-down control from the frontal cortex due to altered signal pathways involving the amygdala and ventral striatum. The serotonergic system is known to be involved in neurodevelopment and neuroplasticity. Therefore, the aim of this study was to investigate whether structural differences in white matter can explain the differences in risk-seeking behaviour that we observed in our previous study and whether *5-HTTLPR* groups differ in their white matter microstructure. These differences can be detected using diffusion-weighted imaging (DWI). We assumed lower structural connectivity in S/S compared to L/L carrier, and a negative relationship between risk-seeking for losses and connectivity. We used DWI to compute diffusion parameters for the frontostriatal and uncinate tract in 175 individuals (39 S/S, 80 S/L, 56 L/L). Results showed no significant relationship between diffusion parameters and risk-seeking for losses. Furthermore, we did not find significant differences in diffusion parameters of the S/S vs. L/L group. There were only group differences in the frontostriatal tract showing stronger structural connectivity in the S/L group, which is also reflected in the whole brain approach. Therefore, the data do not support our hypothesis that the association between *5- HTTLPR* and risk-seeking for losses is related to differences in white matter pathways implicated in decision-making.

## Introduction

Understanding the neurobiological basis of decision-making under risk is an important target in the neuroeconomic community. Kahneman and Tversky (1979) showed with their prospect theory that individuals do not behave rationally when making choices involving risks, but instead show a pattern termed the *reflection effect*. It describes the observation that, when offered a smaller but certain amount of money and a larger but probabilistic amount to gain, individuals are risk-averse, i.e. they prefer the safe option. The opposite pattern, hence *reflection effect*, is shown when the offers are about losing money, either a certain smaller amount or a larger but probabilistic amount. Here individuals usually behave more risk-seeking, i.e. they choose the probabilistic offer more often than the safe option.

This reflection effect has been suspected to stem from emotional responses that bias choices to be more risk-averse or risk-seeking. Indeed, using a risky choice paradigm where offers were framed either as gains or losses, De Martino, Kumaran et al. (2006) found heightened amygdala activation when choices were made in agreement with the reflection effect (sure option in the gain domain, risky option in the loss domain) compared to the opposite behaviour. Further evidence from a mixed gambles task showed that the ventral striatum (VS) codes for the expected value of probabilistic choices in the gain domain and the amygdala for expected value in the loss domain (Yacubian, Gläscher et al. 2006). In general, the striatum and medial parts of the frontal cortex are known to carry valuation signals by integrating outcome related information such as magnitude, probability and delay (for reviews see Rangel, Camerer et al. 2008, Peters and Büchel 2011) and some evidence suggests that the valuation process may be influenced by the amygdala (Gottfried, O’Doherty et al. 2003). Further evidence for a role of the amygdala in loss related decision-making is provided by De Martino, Camerer et al. (2010) who showed that the amygdala mediates loss aversion behaviour possibly by computing an arousal signal related to the prospective monetary loss. Moreover, the amygdala has strong connections to frontal brain regions and recent studies show that its activity is regulated via inhibitory top-down control by the ventromedial prefrontal cortex (vmPFC) and a loss of control can result in potentiated amygdala activity (Motzkin, Philippi et al. 2015). Therefore, the question arises whether the inhibitory top-down control is reflected in the structural makeup of fibre bundles connecting the vmPFC with brain regions such as the amygdala and VS.

Previous research suggests that the *5-HTTLPR*, a natural occurring genetic variation in the promoter region of the gene (SLC6A4) coding for the serotonin transporter (5-HTT), influences neuronal signalling between the amygdala, striatum and vmPFC/OFC (Hariri and Holmes 2006). The 5-HTT is responsible for the reuptake of serotonin (5-HT) from the synaptic cleft to the presynaptic nerve terminal and hence regulates extracellular 5-HT levels. 5-HTT availability and regulatory activity is also very important during the development of the central nervous system as 5-HT influences neuronal plasticity, proliferation and differentiation (for a review see Persico, Kalueff et al. 2010). *5-HTTLPR* regulates the transcriptional efficiency (and hence transporter availability) in a way that one allelic variant with 14 repeats (short or S-allele) results in lower transcriptional efficiency and the other variant with 16 repeats (long or L-allele) results in higher transcriptional efficiency and, eventually, 5-HTT availability. PET imaging with the radioligand [^11^C]DASB revealed a high density of 5-HTT in the striatum, moderate to high in the amygdala and moderate in the vmPFC (Kranz, Kasper et al. 2010, Kobiella, Reimold et al. 2011).

The functional relevance of this genotype was demonstrated in a study by Roiser, de Martino et al. (2009), who used a risky choice fMRI task with a gain and loss frame for the presented offers in participants either homozygous for the L_a_-allele or the S-allele of the *5-HTTLPR* and tested for differences in choice behaviour and brain activation between the genotype groups. Although they did not find significant differences in choice behaviour, they observed increased amygdala activation only in the S/S group when they chose according to the frame effect compared to when they chose contrary to it. The L_a_/L_a_ group did not show such an effect. An additional functional connectivity analysis using a psycho-physiological interaction analysis (PPI) revealed an increased amygdala-PFC coupling only in the L_a_/L_a_ group when making choices counter to the frame effect. Increased amygdala activation, reduced amygdala-PFC coupling and reduced amygdala grey matter volume in S-allele carrier compared to L/L individuals have also been shown in the context of negative emotional stimuli (Pezawas, Meyer-Lindenberg et al. 2005, Kobiella, Reimold et al. 2011). These findings indicate an important functional role for the amygdala in decision-making and emotion processing. Therefore, the reduced functional coupling and concurrent increase in amygdala activation observed in S-allele carrier may be due to a compromised cortico-amygdala pathway (Hariri and Holmes 2006).

In support of this interpretation, a study by Klucken, Schweckendiek et al. (2015) used DTI to investigate white matter microstructural properties of the uncinate tract, a fibre bundle that connects the temporal lobe (including the amygdala) with the inferior frontal lobe (i.e., the vmPFC) together with a fear conditioning paradigm to measure amygdala reactivity across *5- HTTLPR* groups in 107 participants. They found increased amygdala activation in S-allele carrier compared to the L/L group as well as increased fractional anisotropy, a measure of structural connectivity, in the S-allele participants, which they interpreted as elevated bottom-up control. There are, however, two other studies (Pacheco, Beevers et al. 2009, Jonassen, Endestad et al. 2012) that found the opposite result, i.e. reduced fractional anisotropy for S-allele carrier. It should be noted that the former study only tested 33 and the latter 37 females, which makes it difficult to draw a strong conclusion based on their findings.

Another important bundle that has been of high interest in the realm of decision-making research is the frontostriatal (also termed accumbofrontal) tract, which connects the VS with the vmPFC (Rigoard, Buffenoir et al. 2011). Especially in the delay discounting domain, several studies reported a negative relationship between FA values of this tract and temporal discounting rates in young adults (Peper, Mandl et al. 2013) and developing populations in the age range of 8-25 years (Olson, Collins et al. 2009, Achterberg, Peper et al. 2016). These studies suggest a relationship between the structural properties of the frontostriatal tract and delay discounting. However, little is known about how 5-HTTLPR modulates the structural properties of this tract and the relationship of this modulation with probabilistic choice.

In an earlier study, we found a significant relationship between *5-HTTLPR* and risk-seeking for losses (but not for risk-aversion for gains). Specifically, individuals homozygous for the S-allele were more risk-seeking compared to heterozygous and individuals homozygous for the L-allele. This effect was not present when we acutely manipulated 5-HT levels (Neukam, Kroemer et al. 2018) and we speculated that *5-HTTLPR* related differences in white matter structure may account for our finding.

Therefore, the aim of this study is to investigate whether our gene-behaviour association can be explained with differences in individual white matter connectivity. To this end, based on the literature, we focused on the uncinate and frontostriatal fasciculus as a priori volumes of interest as they are most likely to be involved in modulating decision-making and be modulated by *5- HTTLPR*. We assumed increased risk-seeking for losses in S/S carrier to be driven by reduced top-down control from the vmPFC resulting in a stronger emotional response towards losses and, possibly, altered evaluation processes in the striatum. Therefore, we expect reduced structural connectivity of the uncinate and frontostriatal tract in S/S compared to L/L genotype individuals and a negative relationship between the structural connectivity (indicated by higher FA, AD and lower MD, RD) and risk-seeking for losses scores. Finally, we use an exploratory voxel-based approach of the whole brain white matter to investigate possible relationships between other fibre bundles, genotypes and behaviour.

## Methods

### Participants

This study is part of two larger projects that investigated the role of dopamine and serotonin on meta-control parameters and brain function (Neukam, Kroemer et al. 2018, Deza-Araujo, Neukam et al. 2019, Kroemer, Lee et al. 2019). The recruitment was done via standardized invitation letters sent to addresses based on a random sample stratified by sex and age (20-40 years), which were provided by the residential registry. All interested individuals were screened and excluded if one of the following criteria applied: pregnancy; not fulfilling the common criteria for MR safety; a current somatic disease requiring medical treatment; any psychiatric disorders that required pharmacological treatment within the last year; and a lifetime history of one of the following conditions (for ICD-10): organic psychiatric disorders (F0), opiate, cocaine, stimulants, hallucinogens, inhalants or poly-substance dependence, schizophrenia or related personality disorders (F2), and affective disorders (F3). Participants who passed the screening were invited to the study; their visual acuity was checked to ensure that it was at least 0.8. In total 611 participants completed a baseline visit, in which blood was taken to be genotyped for the *5-HTTLPR* and stored at −81°C until further processing. Risk-seeking for losses was measured with a probability discounting for losses task using the value-based decision-making (VBDM) battery (Pooseh, Bernhardt et al. 2018). Afterwards, participants were re-invited to take part either in the dopamine or serotonin project during which diffusion weighted images were acquired. The local ethics review board of the Technische Universität Dresden approved of the study protocols and all participants gave written informed consent in line with the Declaration of Helsinki.

### Probability discounting for losses (PDL) task

In this task, participants had to choose between two offers: a smaller certain loss or a larger probabilistic loss, both simultaneously presented. All offers presented were randomly shown on the left or right side on the computer screen, and the chosen offer was indicated with a red frame. Participants were informed beforehand that one of their choices in every task would be selected randomly and deducted from the total balance they could accumulate during the baseline visit. During the task, they did not get feedback about the outcomes of each choice.

Based on individual choices, the discounting rate (*k*) was estimated assuming hyperbolic discounting, following the formula:

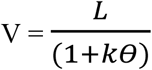

where ϴ = (1 - P)/P is the transformation of reward probability P (2/3, 1/2, 1/3, 1/4, and 1/5) to odds against winning. The loss, L, ranged from −5 Euro to −20 Euro. The task consisted of 30 trials. In all tasks, the likelihood of choosing between the two offers follows a softmax probability function in which β>0 serves as a consistency parameter such that its large values correspond to a high probability of taking the most valuable action. The algorithm starts from liberal prior distributions on the parameters and, after observing a choice at each trial, updates the belief about the parameters using the Bayes’ rule P(k, β | choice) ∞ P(choice| k, β)P(k, β) to find offers close at the individual indifference point. The estimated *k* parameter from the final trial best explains choice behaviour with high *k* values indicating increased risk-seeking as higher but probabilistic losses are discounted and hence preferred over smaller, certain losses. A detailed description of the mathematical framework is reported in Pooseh, Bernhardt et al. (2018). All tasks were implemented in MATLAB (Release 2010a, The MathWorks, Inc., Natick, MA, USA) and the Psychtoolbox 3.0.10 based on the Psychophysics Toolbox extensions (Brainard 1997, Pelli 1997).

### Genotyping

The collected blood samples were sent to the Central Institute of Mental Health in Mannheim, Germany, to perform the genotyping for the *5-HTTLPR*. The exact procedure is described elsewhere (Dukal, Frank et al. 2015). Due to the failure to take blood from nine participants, blood samples from 602 participants were available for genotyping. The observed allele frequency was 39.9% for S and 60.1% for L, with the following genotype groups: 99 S/S, 283 S/L, and 220 L/L. The allele and genotype frequencies did not deviate significantly from the Hardy-Weinberg equilibrium (χ^2^ = 0. 2461, df = 1, p = 0.62).

### Imaging

Diffusion-weighted images (DWI) were acquired on a 3 Tesla Magnetom TrioTim scanner (Siemens Healthcare GmbH, Erlangen, Germany), equipped with a 32-channel head coil. In total 36 transverse scans, consisting of 4 non diffusion-weighted (B0) and 32 non-collinear diffusion-weighted images with a b1000 s/mm^2^ factor, were obtained. Parallel imaging was realized with a GRAPPA factor = 2 while the other parameters were as follows: repetition time (TR) = 9200ms; echo time (TE) = 92 ms; a basis resolution of 128×128×72 mm^3^ with 2.1 mm isotropic voxels (no gap); field-of-view (FOV) = 275×275 mm. Additionally, a high-resolution T1-weighted magnetization prepared rapid acquisition gradient echo (MP-RAGE) image for normalization, anatomical localization as well as screening for structural abnormalities by a neuroradiologist (TR: 1900 ms; TE: 2.26 ms; flip angle: 9°; FOV: 256 × 256 mm; 176 sagittal slices; voxel size: 1 × 1 × 1 mm) was acquired. Preprocessing of the DWI data included motion and eddy current correction using FSL eddy (version 5.0.11) (Andersson and Sotiropoulos 2016), as well as the *-repol* setting to detect and replace slices that can be considered as outliers using default parameter (Andersson, Graham et al. 2016) using the in-house developed NICePype software (Müller, Küttner et al. 2015). All 221 volumes were visually inspected for artefacts. In total, 38 data sets had to be excluded: 26 because of strong absolute rotation ≥ 1° along the x, y, or z axis, 2 showed abnormally large ventricles, and 10 suffered from severe distortion artefacts. The preprocessed images were then loaded into the ExploreDTI toolbox (Leemans, Jeurissen et al. 2009) and the diffusion tensor model was fitted to the data using the RESTORE algorithm (Chang, Jones et al. 2005). Afterwards, the DT matrices of interest were computed: FA, mean diffusivity (MD), axial diffusivity (AD) and radial diffusivity (RD) and deterministic whole brain tractography (Basser, Pajevic et al. 2000) was performed (minimum FA = 0.2, minimal fibre length = 30 mm, maximal fibre length = 300 mm, maximum angle = 30°, cubic interpolation).

### Frontostriatal and uncinate fiber tract selection and volume of interest (VOI) generation

#### Uncinate fasciculus

To obtain a VOI for the uncinate fasciculus, we employed a similar method to the one reported by Schaeffer, Krafft et al. (2014). To this end, we used the TBSS pipeline to warp all individual DTI images to MNI space (FMRIB58 template). The warped FA images were thresholded to include only voxels with a value of at least 0.2 or above. We took the uncinate fasciculus from the John Hopkins University white matter tractography probability atlas (Hua, Zhang et al. 2008) and thresholded the probabilities to greater or equal 5% to exclude less likely voxels. Finally, each normalized and thresholded FA image was combined with thresholded probabilistic tract map and the mean FA skeleton computed during the TBSS procedures described above. The resulting VOIs were then retransformed to native space and applied to the DTI images to extract the parameters of interest. The VOI is shown in Fig. 1(a).

**Fig. 1.**
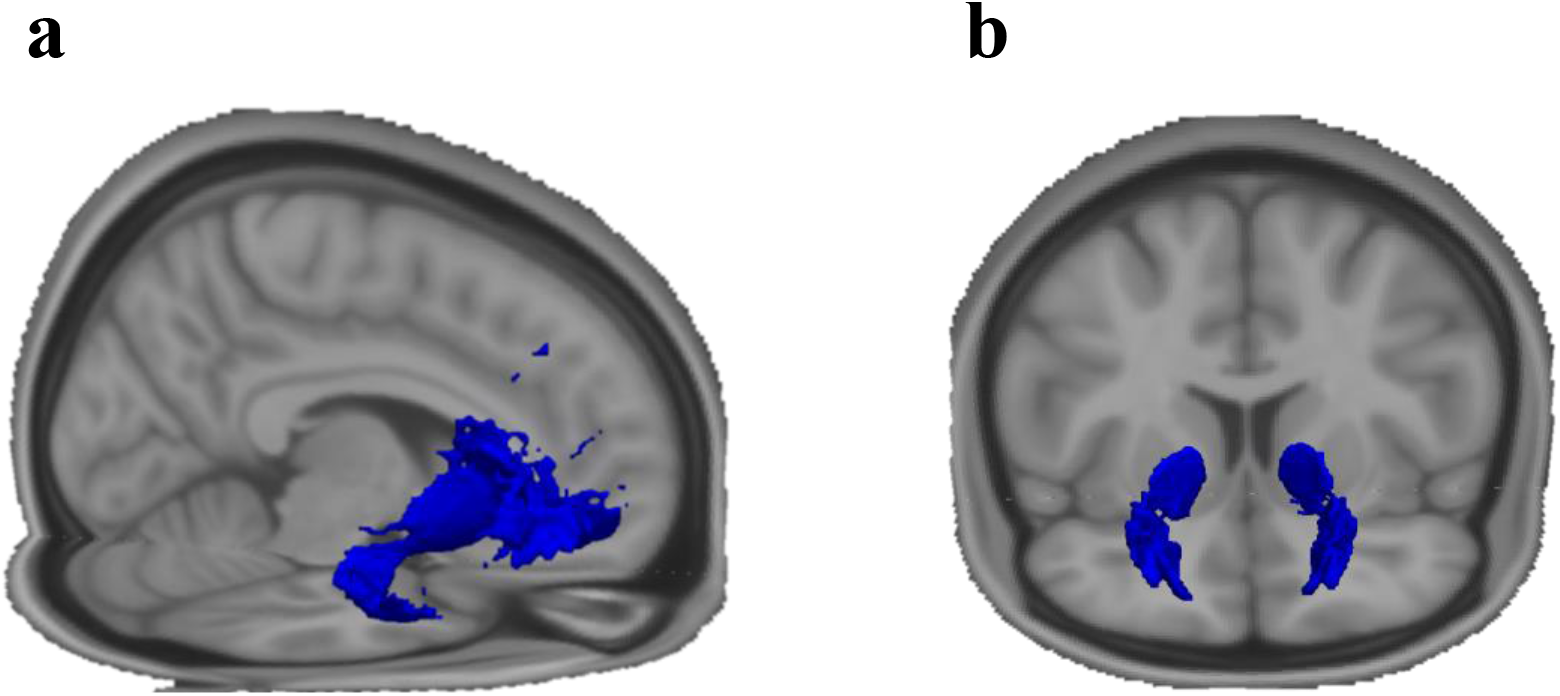
Volumes of interest of the (a) uncinate fasciculus and (b) frontostriatal fasciculus. Both volumes were created in standard space where the uncinate tract was created based on an atlas template and the frontostriatal tract was created based on individual tractography. Only voxels in deep white matter were analyzed based on the FA skeleton created with TBSS. See section 4.2.5. for details.

#### Frontostriatal tract

As the frontostriatal tract is not yet part of white matter atlases, we used a region of interest (ROI) approach, combined with the individually computed tractograms. First, we generated a ROI of the striatum by combining the accumbens, putamen and caudate parts from the Harvard-Oxford subcortical atlas implemented in FSL. Next, we used regions (Frontal_Sup_Orb, Frontal_Med_Orb, Rectus) from the automated anatomic labeling atlas (Tzourio-Mazoyer, Landeau et al. 2002) to create a vmPFC/OFC mask. In order to transform the ROIs from MNI standard space to individual diffusion space, a series of computations were performed. The first step was to register the individual T1 image to MNI space using the *flirt* and *fnirt* algorithms thereby obtaining the non-linear transformation coefficients of interest. The next aim was to average and skull-strip the B0 images and register the averaged image to the individual T1 image using *flirt*. The obtained transformation matrix was then used with *applywarp*, together with the non-linear transformation coefficients to warp the B0 image to MNI space. As we were interested in having the inverse matrices to transform the ROIs from MNI to diffusion space, we used to *convert_xfm* and *invwarp* operations to invert the transformations from MNI to T1 space and from T1 to diffusion space. The inverted matrices and the *applywarp* command were then used to transform the striatum and vmPFC/OFC masks from MNI to diffusion space.

The individual tractograms and transformed ROIs were next loaded into TrackVis (version 0.6.1; Wang and Wedeen 2015) and streamlines that pass from the striatum to the vmPFC/OFC or vice versa were generated. Exclusion masks were individually set on the mid sagittal plane, on the coronal plane at the splenium of the corpus callosum, and on the axial plane on the level of the anterior temporal gyrus. If necessary, single spurious streamlines were additionally manually removed. The resulting tracts were then exported as nifti files.

A prerequisite for the creation of a group template was the employment of the FSL tract-based spatial statistics (TBSS) pipeline (Smith, Jenkinson et al. 2006), which warps individual FA maps to MNI space (the FMRIB58 template provided by FSL), averages all FA images to compute a mean FA image, which is then reduced to a skeleton, based on voxels from the nearest tract centre. In a next step, all tracts were non-linearly warped to the MNI template, registered to the FA skeleton using the *tbss_non_fa* command from TBSS and binarized. Finally, all tracts were summed up into one nifti, normalized by the number of participants, and thresholded to contain only voxels that exist in at least 50% of the sample, and binarized again. This group VOI was then retransformed to native space and applied to the DTI metrics of interest (FA, MD, AD, RD) to extract the tract related metrics. The VOI is depicted in Fig. 1(b).

### ROI statistical analysis

To address the question whether there is a relationship between *5-HTTLPR* groups, white matter structure and risk-seeking for losses, we used a multivariate analysis of covariance (MANCOVA) for each tract with FA, AD, MD and RD as dependent variables with genotype (S/S, S/L, L/L) as group factor and logarithm of *k* from the PDL task as covariate of interest, as well as sex and age as control variables. We set our statistical threshold of significance at p < .05.

### Whole brain analysis

To explore potential effects of *5-HTTLPR* and risk-seeking for losses in other regions of the brain, we used the TBSS pipeline described above to skeletonize all FA, MD, AD, and RD images. We used voxelwise non-parametric statistical analyses based on 10.000 random permutations and the threshold-free cluster enhancement (TFCE) approach to test for the main effect of genotype (S/S, S/L, L/L), the main effect of risk-seeking for losses and interaction effects while controlling for sex and age. Additionally, age and sex were demeaned and entered as covariate regressors as they were found in previous studies to be related to the DTI parameters (Menzler, Belke et al. 2011, Jonassen, Endestad et al. 2012). We assumed significance at a family-wise error corrected p-value of < .05. Classification of tracts the clusters belong to was performed with the JHU White Matter Tractography Atlas (Hua, Zhang et al. 2008). To further explore the contribution of each genotype to all significant clusters, binarized masks were generated from them and the four DTI parameters were extracted and averaged across cluster voxels, separately for each parameter. In a next step, a multivariate ANOVA was conducted with the four DTI parameters as dependent variables and genotype as predictor.

## Results

### Sample information

Eight participants out of the 183 had missing PDL data resulting in 175 data sets for analysis. This sample consisted of 39 (14 females) S/S carrier, 56 (12 females) S/L carrier and 80 (35 females) L/L carrier demonstrating an overall number of males than females (χ^2^ = 7.252, df = 2, p = .027). Age was similarly distributed among genotype groups (F_2,169_ = 3.027, p = .051): S/S (males: 32.8 ± 6.1, females 31.4 ± 5.5 years), S/L (males: 34.8 ± 4.3, females 34.9 ± 4.7 years), S/L (males: 32.8 ± 6.1, females 31.4 ± 5.5 years), and L/L (males: 33.8 ± 5.1, females 32.3 ± 5.9 years).

### Tract analysis

To test our first hypothesis, we used a multivariate analysis of variance (MANOVA) separately for each tract and investigated simple linear contrasts to compare the homozygous S- and L-allele groups, controlling for sex and age. For the second hypothesis, we used partial Pearson correlations to investigate the relationship between the *k* values (on log scale to approximate a normal distribution) from PDL and the DTI parameters, controlling for *5-HTTLPR*, sex and age.

#### Association between 5-HTTLPR and the frontostriatal tract

The simple linear contrast analyses between S/S and L/L groups did not reveal any significant differences for each of the DTI metrics (all p > .06). See Table 1 and Fig. 2 for details. The results did also not change when we combined the S/S and S/L group.

**Table 1.**
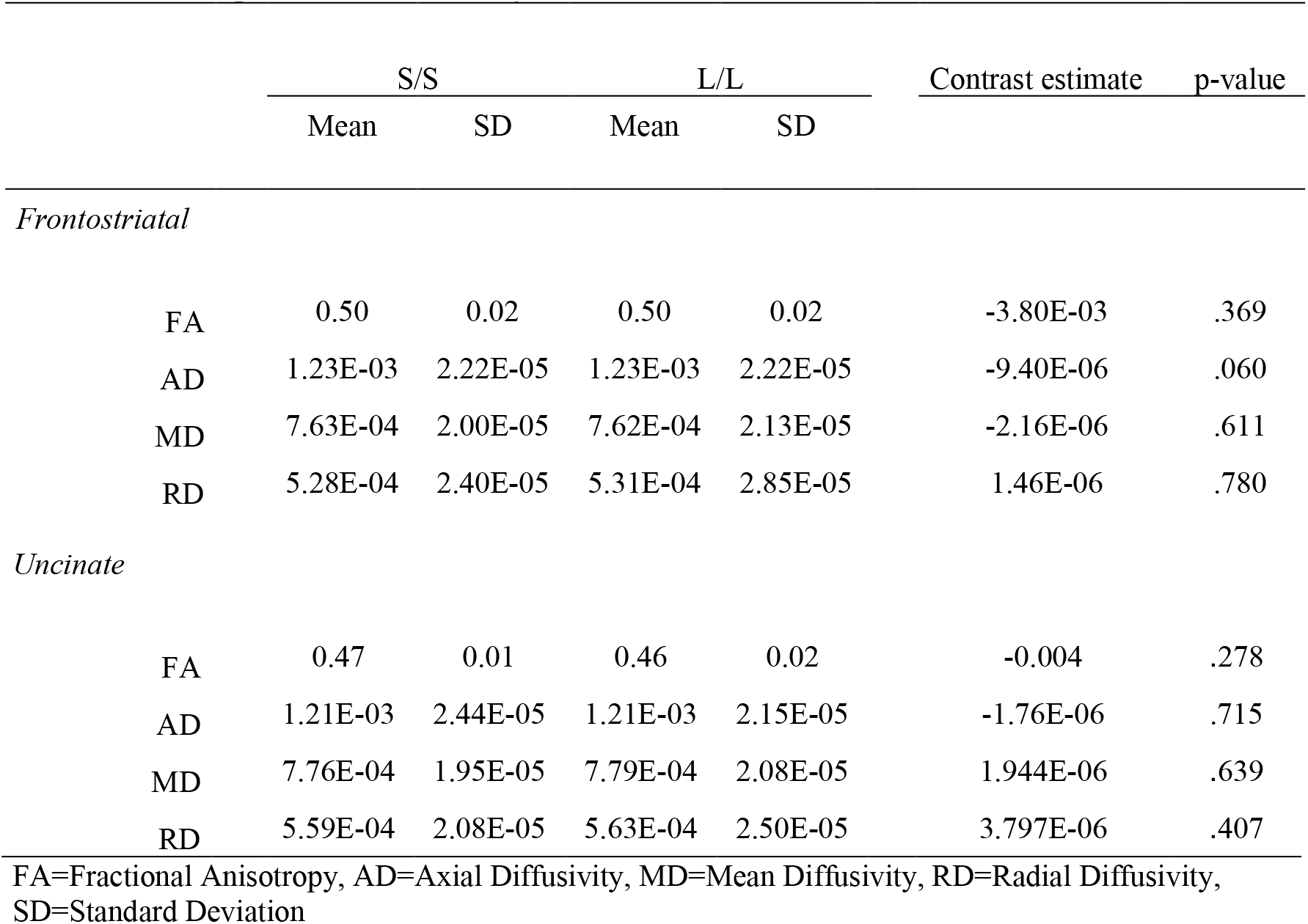
Results of the simple linear contrast analyses

**Fig. 2.**
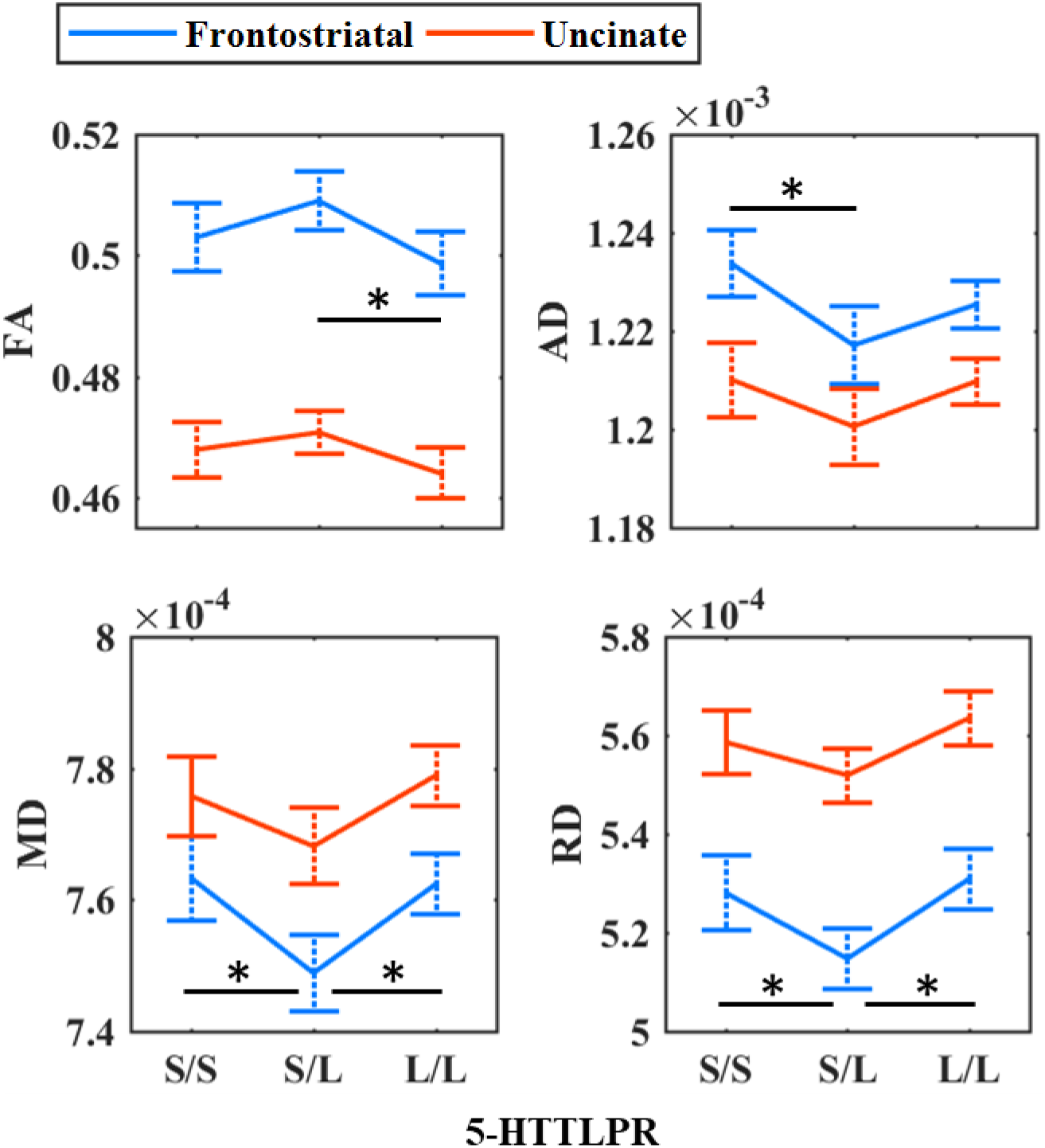
Main effect of *5-HTTLPR* on DTI parameters: fractional anisotropy (FA), axial diffusivity (AD), mean diffusivity (MD) and radial diffusivity (RD). Error bars are bootstrapped with 10.000 iterations and denote 95% bias corrected and accelerated confidence intervals. *p < .05

However, the multivariate analysis showed significant main effects of genotype (Wilk’s Λ = 0.924, F_6,330.000_ = 2.222, p = .041, ηp^2^ = .039), sex (Wilk’s Λ = 0.953, F_3,165.000_ = 2.689, p = .048, ηp^2^ = .047) and age (Wilk’s Λ = 0.952, F_6,330.000_ = 2.222, p = .041, ηp^2^ = .039). For the main effect of genotype, an exploratory oneway ANOVA with Games-Howell post-hoc tests showed that S/L individuals had significantly higher FA values than L/L individuals (p = .013), S/S had higher AD values compared to S/L individuals (p = .007), while S/S and L/L carrier had higher MD (p_S/S_ = .004; P_L/L_ = .002) and RD values (p_S/S_ = .024; p_L/L_ = .001) compared to S/L carrier. Furthermore, post-hoc independent sample t-tests demonstrated that the effect of sex was related to higher FA (t_173_ = −2.529, p = .012) lower MD (t_173_ = 2.962, p = .003) and RD (t_173_ = 2.888, p = .004), but not AD (t_173_ = 1.593, p = .113) in males compared to females. Finally, exploratory Pearson’s correlation showed that age was significantly negatively correlated with AD (r = −.219, p = .004), but not with FA (r = .005, p = .948) nor MD (r = −.143, p = .059) nor RD (r = −.073, p = .336). Descriptive statistics are shown in Table 2.

**Table 2.**
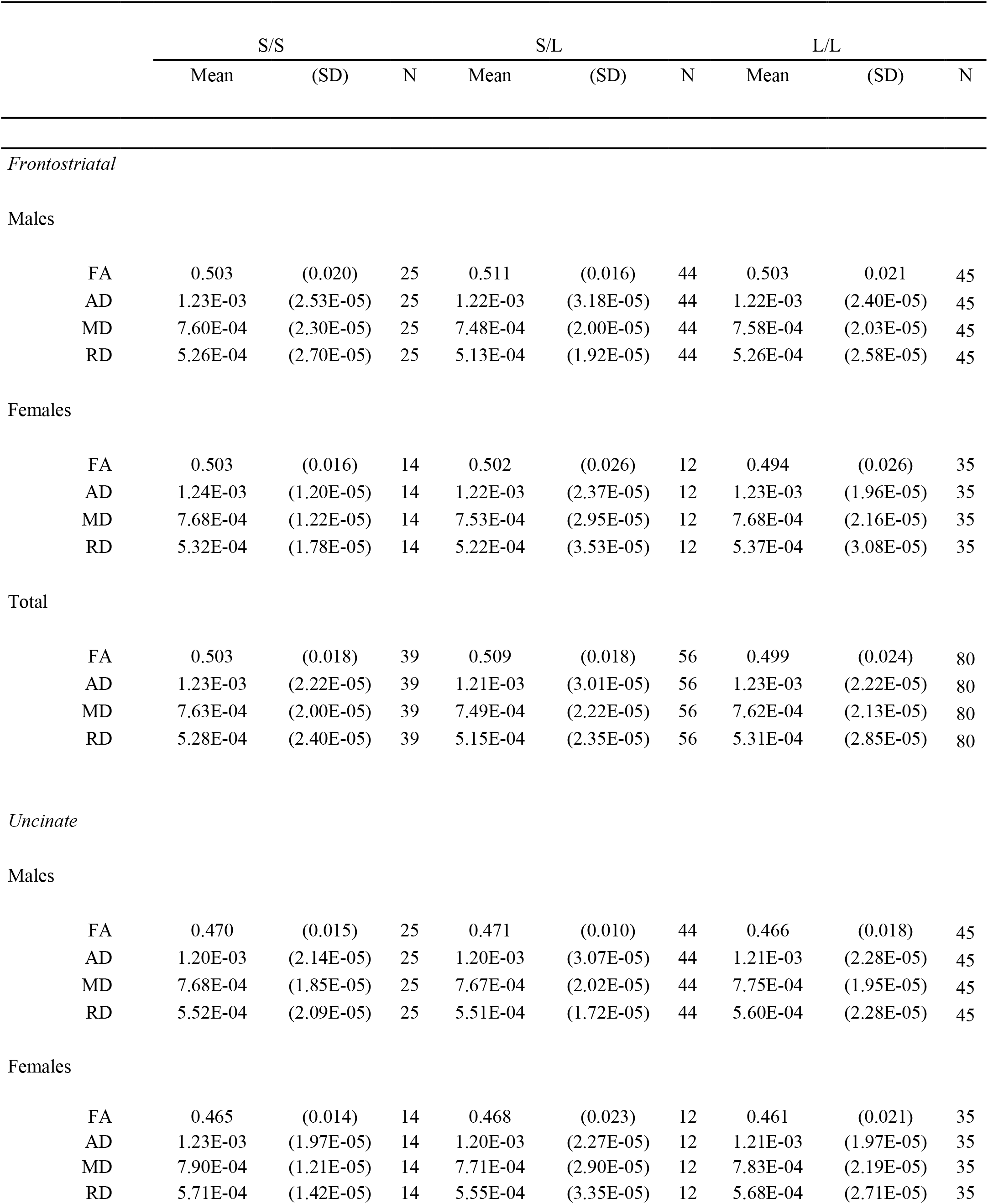
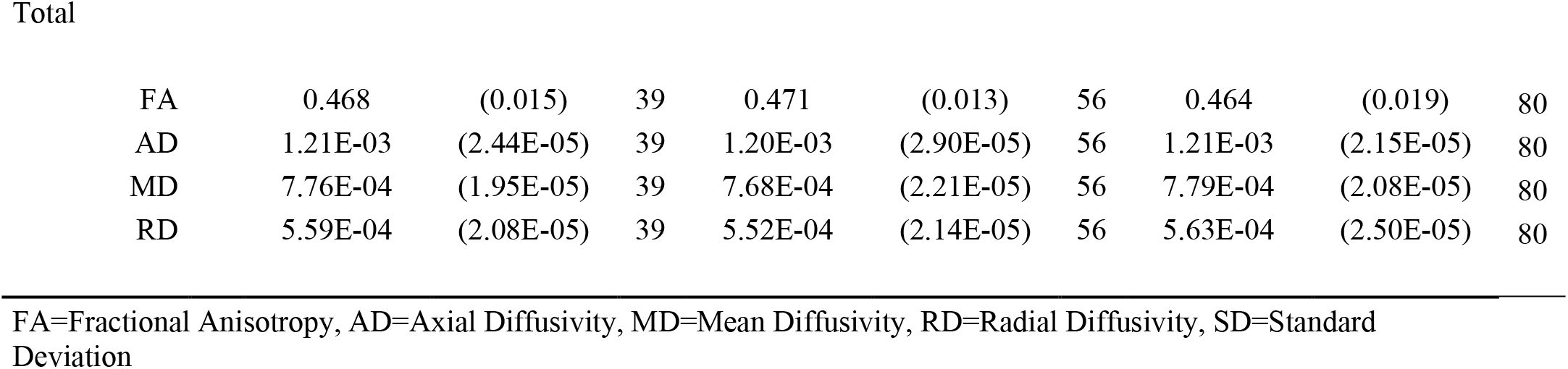
Associations between *5-HTTLPR* and the frontostriatal/uncinate tract

|  |  | S/S |  |  | S/L |  |  | L/L |  |  |
| --- | --- | --- | --- | --- | --- | --- | --- | --- | --- | --- |
|  |  | Mean | (SD) | N | Mean | (SD) | N | Mean | (SD) | N |
| *Frontostriatal* |  |  |  |  |  |  |  |  |  |  |
| Males |  |  |  |  |  |  |  |  |  |  |
|  | FA | 0.503 | (0.020) | 25 | 0.511 | (0.016) | 44 | 0.503 | 0.021 | 45 |
|  | AD | 1.23E-03 | (2.53E-05) | 25 | 1.22E-03 | (3.18E-05) | 44 | 1.22E-03 | (2.40E-05) | 45 |
|  | MD | 7.60E-04 | (2.30E-05) | 25 | 7.48E-04 | (2.00E-05) | 44 | 7.58E-04 | (2.03E-05) | 45 |
|  | RD | 5.26E-04 | (2.70E-05) | 25 | 5.13E-04 | (1.92E-05) | 44 | 5.26E-04 | (2.58E-05) | 45 |
| Females |  |  |  |  |  |  |  |  |  |  |
|  | FA | 0.503 | (0.016) | 14 | 0.502 | (0.026) | 12 | 0.494 | (0.026) | 35 |
|  | AD | 1.24E-03 | (1.20E-05) | 14 | 1.22E-03 | (2.37E-05) | 12 | 1.23E-03 | (1.96E-05) | 35 |
|  | MD | 7.68E-04 | (1.22E-05) | 14 | 7.53E-04 | (2.95E-05) | 12 | 7.68E-04 | (2.16E-05) | 35 |
|  | RD | 5.32E-04 | (1.78E-05) | 14 | 5.22E-04 | (3.53E-05) | 12 | 5.37E-04 | (3.08E-05) | 35 |
| Total |  |  |  |  |  |  |  |  |  |  |
|  | FA | 0.503 | (0.018) | 39 | 0.509 | (0.018) | 56 | 0.499 | (0.024) | 80 |
|  | AD | 1.23E-03 | (2.22E-05) | 39 | 1.21E-03 | (3.01E-05) | 56 | 1.23E-03 | (2.22E-05) | 80 |
|  | MD | 7.63E-04 | (2.00E-05) | 39 | 7.49E-04 | (2.22E-05) | 56 | 7.62E-04 | (2.13E-05) | 80 |
|  | RD | 5.28E-04 | (2.40E-05) | 39 | 5.15E-04 | (2.35E-05) | 56 | 5.31E-04 | (2.85E-05) | 80 |
| *Uncinate* |  |  |  |  |  |  |  |  |  |  |
| Males |  |  |  |  |  |  |  |  |  |  |
|  | FA | 0.470 | (0.015) | 25 | 0.471 | (0.010) | 44 | 0.466 | (0.018) | 45 |
|  | AD | 1.20E-03 | (2.14E-05) | 25 | 1.20E-03 | (3.07E-05) | 44 | 1.21E-03 | (2.28E-05) | 45 |
|  | MD | 7.68E-04 | (1.85E-05) | 25 | 7.67E-04 | (2.02E-05) | 44 | 7.75E-04 | (1.95E-05) | 45 |
|  | RD | 5.52E-04 | (2.09E-05) | 25 | 5.51E-04 | (1.72E-05) | 44 | 5.60E-04 | (2.28E-05) | 45 |
| Females |  |  |  |  |  |  |  |  |  |  |
|  | FA | 0.465 | (0.014) | 14 | 0.468 | (0.023) | 12 | 0.461 | (0.021) | 35 |
|  | AD | 1.23E-03 | (1.97E-05) | 14 | 1.20E-03 | (2.27E-05) | 12 | 1.21E-03 | (1.97E-05) | 35 |
|  | MD | 7.90E-04 | (1.21E-05) | 14 | 7.71E-04 | (2.90E-05) | 12 | 7.83E-04 | (2.19E-05) | 35 |
|  | RD | 5.71E-04 | (1.42E-05) | 14 | 5.55E-04 | (3.35E-05) | 12 | 5.68E-04 | (2.71E-05) | 35 |

**Table 2 (cont.)** Total
|  |  |  |  |  |  |  |  |  |  |
| --- | --- | --- | --- | --- | --- | --- | --- | --- | --- |
| FA | 0.468 | (0.015) | 39 | 0.471 | (0.013) | 56 | 0.464 | (0.019) | 80 |
| AD | 1.21E-03 | (2.44E-05) | 39 | 1.20E-03 | (2.90E-05) | 56 | 1.21E-03 | (2.15E-05) | 80 |
| MD | 7.76E-04 | (1.95E-05) | 39 | 7.68E-04 | (2.21E-05) | 56 | 7.79E-04 | (2.08E-05) | 80 |
| RD | 5.59E-04 | (2.08E-05) | 39 | 5.52E-04 | (2.14E-05) | 56 | 5.63E-04 | (2.50E-05) | 80 |

FA=Fractional Anisotropy, AD=Axial Diffusivity, MD=Mean Diffusivity, RD=Radial Diffusivity, SD=Standard Deviation

#### Association between 5-HTTLPR and the uncinate tract

There was no significant difference between the two homozygous groups with respect to the DTI parameters (all p > .28). Results are also shown in Table 1 and Fig. 2. Similarly, as above, combining the S-allele groups did not change the results significantly.

For this tract, the multivariate analysis showed no significant main effects of genotype (Wilk’s Λ = 0.958, F_6,330.000_ = 1.192, p = .310, ηp^2^ = .021), but a significant effect of sex (Wilk’s Λ = 0.952, F_3,165.000_ = 2.786, p = .042, ηp^2^ = .048) and age (Wilk’s Λ = 0.938, F_6,330.000_ = 3.643, p = .014, ηp^2^ = .062). Exploratory post-hoc independent sample t-tests demonstrated that the effect of sex was related to a slightly higher FA (t_96.101_ = −2.024, p = .046) lower AD (t_173_ = 2.828, p = .005), MD (t_173_ = 3.550, p < .001) and RD (, t_173_ = 3.315, p = .001) in males compared to females. For the age effect, Pearson’s correlation showed that age was again significantly negatively correlated with AD (r = −.242, p = .001), but not with FA (r = −.064, p = .397) nor MD (r = −.136, p = .073) nor RD (r = −.057, p = .455). Descriptive statistics are depicted in Table 2.

#### Correlations between frontostriatal/uncinate tract and PDL

The results are depicted in Fig. 3. In brief, there were no significant correlations (all p > .14) between DTI parameters of each tract with PDL as shown in Table 3.

**Fig. 3.**
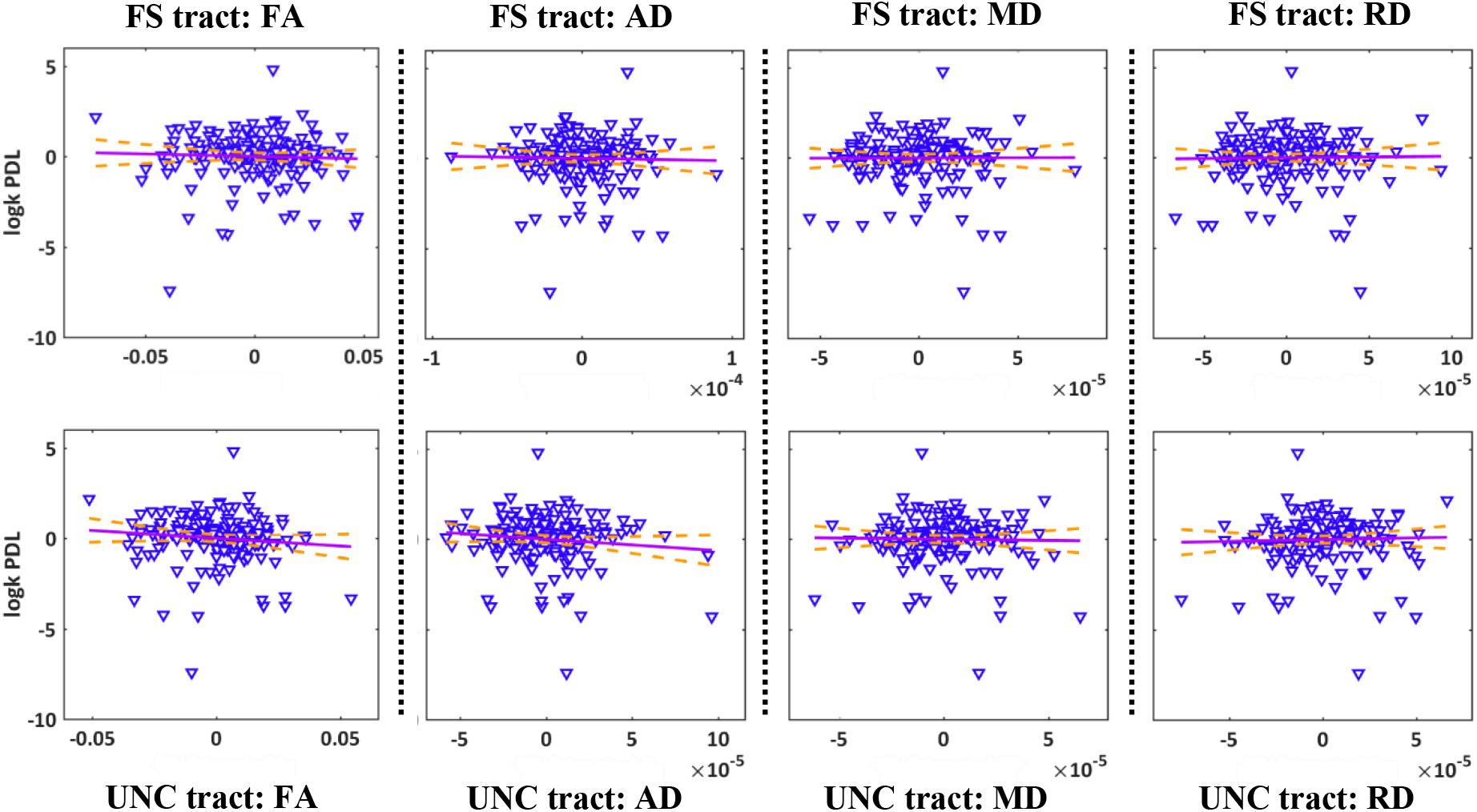
Correlations between risk-seeking for losses (logk PDL) and DTI parameters: fractional anisotropy (FA), axial diffusivity (AD), mean diffusivity (MD) and radial diffusivity (RD). All DTI parameters are unstandardized residuals after controlling for *5-HTTLPR*, sex and age. Dashed lines indicate 95% confidence intervals.

**Table 3.** Correlations of logk PDL with the DTI parameters

| *DTI parameter* | Pearson's r | p-value |
| --- | --- | --- |
| *Frontostriatal tract* |  |  |
| FA | -0.042 | 0.584 |
| AD | -0.024 | 0.757 |
| MD | 0.004 | 0.956 |
| RD | 0.016 | 0.830 |
| *Uncinate tract* |  |  |
| FA | -0.104 | 0.170 |
| AD | -0.111 | 0.142 |
| MD | -0.020 | 0.798 |
| RD | 0.032 | 0.672 |

FA=Fractional Anisotropy, AD=Axial Diffusivity, MD = Mean Diffusivity, RD=Radial Diffusivity

### Whole brain analysis

The results for the main effect of genotype are summarized in Table 4. There were no significant clusters for the main effect of PDL, nor for the interaction with genotype.

**Table 4.** Summary cluster map of the TBSS results for the main effect of genotype

| Summary cluster map of the FDR results for the main effect of genotype |  |  |  |  |  |  |  |
| --- | --- | --- | --- | --- | --- | --- | --- |
| Tracts | Side | Peak voxel (MNI) |  |  | F-Statistic | Cluster size > 100<br>(voxels) | Cluster p-value |
|  |  | x | y | z |  |  |  |
| FA |  |  |  |  |  |  |  |
| SFOF | Left | -22 | -2 | 19 | 19.5 | 10610 | 0.001 |
| ILF | Right | 45 | -11 | -27 | 14.2 | 6452 | 0.003 |
| UNC, IFOF | Right | 18 | 24 | -12 | 13.8 | 669 | 0.015 |
| AD |  |  |  |  |  |  |  |
| Unclassified | Left | -10 | -1 | -14 | 17.3 | 12958 | 0.001 |
| ILF | Right | 40 | -22 | -21 | 11.5 | 1739 | 0.014 |
| Forceps minor | Right | 12 | 31 | 8 | 10 | 1522 | 0.028 |
| UNC, IFOF | Right | 28 | 14 | -10 | 9.84 | 786 | 0.03 |
| Unclassified | Right | 1 | 10 | 14 | 7.64 | 188 | 0.047 |
| Forceps minor | Left | -12 | 29 | -12 | 9.1 | 157 | 0.042 |
| SLF | Left | -34 | -37 | 21 | 10.3 | 141 | 0.038 |
| ATR, IFOF | Right | 23 | 26 | 23 | 5.94 | 141 | 0.047 |
| MD |  |  |  |  |  |  |  |
| ATR | Left | -11 | -17 | -2 | 18.1 | 31113 | 0.001 |
| RD |  |  |  |  |  |  |  |
| ATR | Left | -23 | -2 | 17 | 18.8 | 14709 | 0.001 |
| ILF | Right | 45 | -10 | -28 | 15.3 | 6641 | 0.004 |
| UNC, IFOF | Right | 18 | 24 | -12 | 13.4 | 209 | 0.039 |

FA = Fractional Anisotropy; AD = Axial Diffusivity; MD = Mean Diffusivity; RD = Radial Diffusivity; SFOF = Superior Fronto-Occipital Fasciculus; ILF = Inferior Longitudinal Fasciculus; UNC = Uncinate Fasciculus; SLF = Superior Longitudinal Fasciculus; ATR = Anterior Thalamic Radiation

Simple linear contrast analyses in the context of each of the four MANOVAs showed that S/S individuals had higher FA values (M_S/S_ = 0.48 ± 0.018, M_L/L_ = 0.47 ± 0.017, p = .020, Cohen’s d = −0.434) and lower RD values (M_S/S_ = 5.44×10^-4^ ± 2.21×10^-5^, M_L/L_ = 5.47×10^-4^ ± 2.46×10^-5^, p = .021, Cohen’s d = 0.414) compared to L/L individuals. Visual depictions of the main effect of genotype are shown in the supplemental material Figs. S1/S2.

## Discussion

The purpose of this study was to elucidate whether the association between risk-seeking for losses, as measured with a PDL task, and *5-HTTLPR* we observed earlier (Neukam, Kroemer et al. 2018) may be explained by differences in white matter connecting brain regions that are involved in value-based decision-making (vmPFC, VS, amygdala). Based on existing literature, we chose the frontostriatal and uncinate tract. The former connects the VS with the vmPFC and the latter the vmPFC with the amygdala. Interestingly, the frontostriatal tract has been implicated in decision-making behaviour, but not in *5-HTTLPR*, while the opposite is true for the uncinate tract. This study extends on these findings by examining the association of *5- HTTLPR* with both tracts and furthermore by investigating the relationship between the tracts and risk-seeking for losses, which has not been published before according to the authors knowledge.

The results of all data analyses can be summarized in two parts. First, the DTI parameters (FA, AD, MD, RD) are not related to the discounting rates of the PDL task, neither in the tracts of interest nor with whole brain white matter. Therefore, differences in white matter structure cannot explain risk-seeking for losses in our sample. Second, we did not find the expected linear relationship between genotype and DTI parameters (i.e. higher FA, AD and lower MD, RD for L/L compared to S/S carrier) neither in the frontostriatal, nor in the uncinate tract, nor in other white matter bundles. Hence, we could not replicate previous findings indicating reduced structural connectivity in S/S compared to L/L carrier (Pacheco, Beevers et al. 2009, Jonassen, Endestad et al. 2012).

### White matter and risk-seeking for losses

There are not many reports that studied the contribution of white matter microstructure to value-based decision-making. The majority of existing studies focused on intertemporal choice and found negative correlations between white matter microstructure (such as the frontostriatal tract) and delay discounting (i.e. higher structural connectivity and reduced temporal discounting rates) in longitudinal studies examining participants ranging between 8-26 years (Olson, Collins et al. 2009, Achterberg, Peper et al. 2016) and young adult populations ranging between 18-25 years (Peper, Mandl et al. (2013), van den Bos, Rodriguez et al. (2014); but see Hampton, Alm et al. (2017) for an opposite finding). Much less is specifically known about the relationship between risk-seeking for losses and the uncinate fasciculus. The main motivation to select this tract was that it denotes an important pathway connecting the amygdala to the vmPFC. Research in humans and mice indicated that the frontal cortex regulates the amygdala by reducing its activation in the wake of intense events (Sander, Grafman et al. 2003, Adhikari, Lerner et al. 2015, Motzkin, Philippi et al. 2015). Therefore, reduced structural connectivity may be associated with reduced top-down control, higher amygdala activity and, finally, increased risk-seeking for losses (De Martino, Kumaran et al. 2006).

However, our preliminary findings do not support the conclusion of the studies investigating the frontostriatal tract that higher impulsivity (steeper discounting) is associated with reduced structural connectivity in the context of risk-seeking for losses. It is tempting to speculate that age may be a reason for our null finding as all previous studies had much younger samples. Karlsgodt, John et al. (2015) for example showed that the frontostriatal tract microstructure (i.e. FA) increases steadily during childhood until the early twenties to stabilize and slowly decreases around the age of forty. Hence, we have not been able to capture developmental aspects of the decision-making related white matter, in contrast to the studies above, which may explain our null finding. Still, as there are currently no directly relatable data published it seems premature to draw a final conclusion on whether our finding is a true or false negative.

### *5-HTTLPR* and white matter

Previous studies have shown interest in understanding the white matter microstructure of the uncinate fasciculus in relation to the *5-HTTLPR* because numerous studies using functional and morphometric measures suggest that S-allele carrier have increased amygdala activity (Hariri, Mattay et al. 2002, Heinz, Smolka et al. 2007), reduced grey matter volume (Pezawas, Meyer-Lindenberg et al. 2005, Kobiella, Reimold et al. 2011) and reduced coupling of the amygdala to the frontal cortex (Pezawas, Meyer-Lindenberg et al. 2005) compared to L/L carrier. This is in line with the hypothesis that there is a gene-dose-effect, where the gene function increases with the number of L-alleles (Hu, Lipsky et al. 2006, Wendland, Moya et al. 2008). Such a relationship was found in two studies investigating the uncinate fasciculus that showed increasing FA values with the number of L-alleles (Pacheco, Beevers et al. 2009, Jonassen, Endestad et al. 2012). Due to the observation that S/S and S/L individuals have similar 5-HTT expression rates and also score similarly on behavioural measures such as trauma exposure (Goldman, Glei et al. 2010), neuroticism (Lesch, Bengel et al. 1996) and depressive symptoms (Neumeister, Hu et al. 2006), studies combine S-allele groups (S/S, S/L) and compare them to L/L carrier. Such an approach was conducted by Klucken, Schweckendiek et al. (2015) who found the opposite pattern of FA values (S > L/L), but did not find any association for genotype and FA in a replication study (Klucken, Tapia Leon et al. 2018). This latter finding is in line with our observation that genotype does not significantly affect FA nor AD, MD and RD in the uncinate tract and the fact that Jonassen, Endestad et al. (2012) and Pacheco, Beevers et al. (2009) only analysed 33 and 37 females, respectively, limits the generalizability of their studies. Additionally, Klucken, Schweckendiek et al. (2015) found the opposite in 100 participants containing both sexes in a first study and no genotype effect in their replication study including 114 participants and finally our null finding with 175 participants supports the notion that the genotype effect is either very small or depends on other presently unknown third variables, following the arguments brought forward by Klucken, Tapia Leon et al. (2018).

We also did not find the expected genotype effect, reduced structural connectivity in S/S compared to L/L carrier, in the frontostriatal tract. Instead, we found non-linear effects of genotype in AD, MD and RD demonstrating less MD in S/L compared to S/S and L/L carrier, less AD in S/L compared to S/S carrier and lower RD in S/L compared to L/L carrier. These findings are not intuitive and are at odds with a gene-dose-effect. An explanation may be a larger proportion of individuals showing molecular heterosis in our sample, a phenomenon that describes heterozygosity in a given genetic polymorphism can result either in a greater expression (positive heterosis) or lesser expression (negative heterosis) of a phenotype compared to homozygosity and such observation may occur in up to 50% of all human genetic association studies (Comings and MacMurray 2000). In case of *5-HTTLPR* there is research reporting such findings in the context of 5-HTT binding potential or 5-HTT availability where S/L individuals had lower scores compared to S/S and L/L (Little, McLaughlin et al. 1998, van Dyck, Malison et al. 2004). Furthermore, Malmberg, Wargelius et al. (2008) reported that male S/L adolescents had higher scores for disruptive behavioural disorder and Steffens, Taylor et al. (2008) observed higher white matter volume lesions in geriatric depressed patients in comparison to the homozygous groups. Our results are in line with these observations but the mechanisms behind heterosis are not yet understood. Comings and MacMurray (2000) suggest three possible reasons: the first being an (inverted) U-shape function indicating that both too little or too much expression has adverse consequences and only intermediate expression is advantageous; the second being an independent third factor causing a hidden stratification of the sample such that in one set S/S carrier have the highest/lowest phenotypic expression and in the second set L/L carrier have the highest/lowest phenotypic expression. The third reason may be greater fitness in heterozygous individuals because they show a broader range of gene expression compared to the homozygous groups. Nevertheless, given that this is the first study reporting such a finding with DTI parameters in the frontostriatal tract, more studies are needed to support this finding.

### Limitations

One possible limitation is that our diffusion-weighted imaging sequence was not sensitive enough to find correlations between the DTI parameters and risk-seeking for losses as well as the expected linear relationship with *5-HTTLPR*. However, despite the fact that we only collected data from 32 direction whereas newer sequences acquire data from twice our number or even more directions, we believe that our number of directions is sufficient to estimate the tensor model and, importantly, to replicate an often published finding that males show consistently higher FA and lower MD and RD compared to women in the frontostriatal and uncinate tract (while the results of AD are inconclusive), which is in line with previous findings that males have a higher structural connectivity compared to females in several brain regions (Westerhausen, Kreuder et al. 2004, Menzler, Belke et al. 2011, van Hemmen, Saris et al. 2017). Another limitation is that we could not use the triallelic *5-HTTLPR* model, which may have given us more information about the reliability of the heterosis effect.

## Conclusion

Overall, we did not find a significant correlation between white matter parameter and risk-seeking for losses in two highly relevant fibre bundles, nor the expected association with respect to *5-HTTLPR*, which may have explained the genotype-behaviour finding in our earlier study (Neukam, Kroemer et al. 2018) and supports the idea of reduced top-down control in S/S compared to L/L individuals. Nevertheless, we found some evidence for the potential existence of heterosis in the frontostriatal tract that needs validation from future studies.

## Acknowledgments

We would like to thank, our student assistants and the medical staff for helping with the recruitment process and data collection, and the radiographers at the neuroimaging centre, as well as our colleagues from the dopamine project, especially Ying Lee. Finally, we thank our participants for their time and effort.

## Supplemental Material

**Fig. S1:**
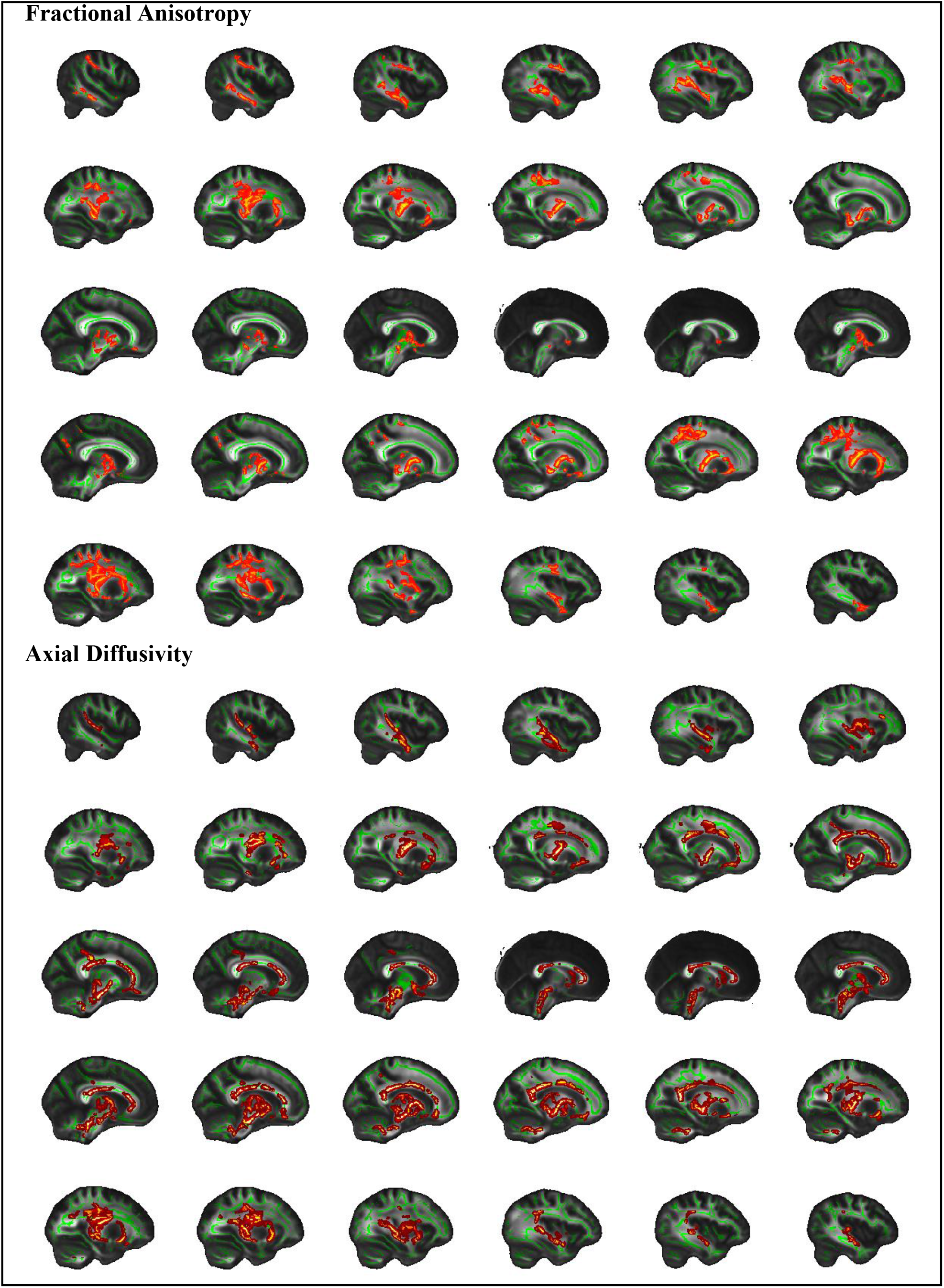
TBSS main effect of *5-HTTLPR* based on 10.000 permutations for fractional anisotropy and axial diffusivity. All voxels survive a family-wise error correction of p < .05.

**Fig. S2:**
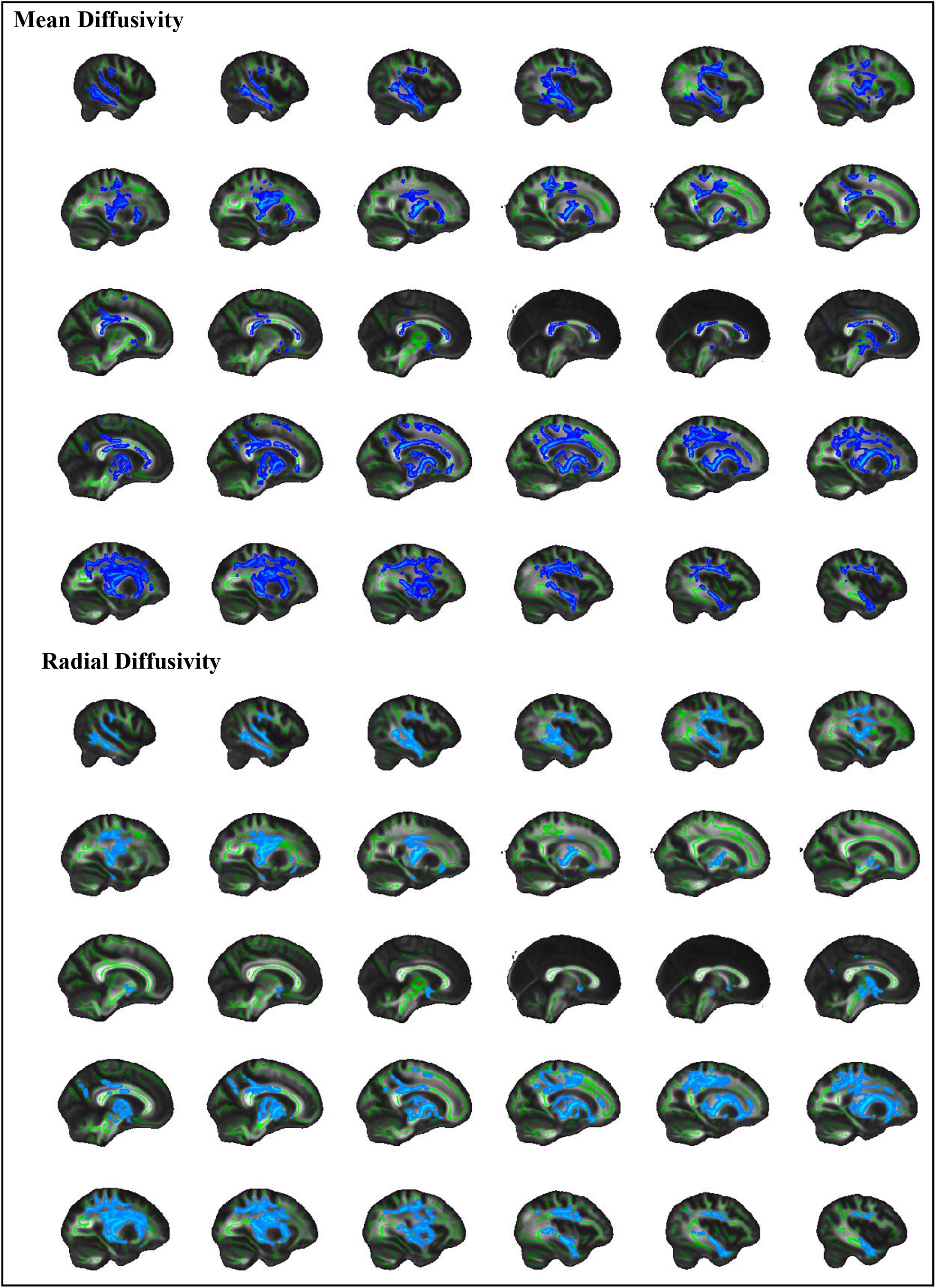
TBSS main effect of *5-HTTLPR* based on 10.000 permutations for mean diffusivity and radial diffusivity. All voxels survive a family-wise error correction of p < .05.

